# What is happening in Dorsal Root Ganglia? A computational analysis of cross-excitation phenomenon

**DOI:** 10.1101/2025.11.11.687782

**Authors:** Dmitrii Perevozniuk, Oleg Gorskii, Pavel Musienko, Nikolay Koshev

## Abstract

The phenomenon of cross-excitation - interaction between neurons of a dorsal root ganglion (DRG) in intact organism, presents a great challenge since its discovery. It can not be addressed to classical neuron-neuron interactions, as the mentioned neurons are believed to be physiologically isolated by glial sheaths from one another. In this work we test existing hypotheses which explain the phenomenon from both biophysical and statistical perspectives. Our conclusion is that neither each proposed mechanism alone, nor interactions of them can adequately describe cross-excitation in normal conditions. We wish to draw greater attention towards the phenomenon, as it seems to be an important part of normal sensory information processing.

**Author summary:** Cross-excitation is a phenomenon discovered in Dorsal Root Ganglia (DRG), whereby activation of one neuron leads to increased excitability of its neighbors. As the mechanism is still poorly understood, we present a computational modelling study, in which we tested primary hypotheses which were proposed to the phenomenon.

The objectives were to test two main ideas - diffusion-mediated and gap-junction mediated cross-excitation in biologically plausible setups.

Firstly, we evaluated a diffusion-based hypothesis, which states that diffusion of K+ ions is the main mechanism by which cross-excitation is mediated. Our results suggest miniscule effect of ion diffusion, even if tightly-packed space of DRG is considered.

Secondly, we analyzed the gap-junction hypothesis, which relates the phenomenon to physical connection between neurons’ glial envelopes. According to the results, this solution would require for each neuron to have an enormous amount of immediate neighbors, in order to sustain high-enough prevalence of connection. Geometrical conclusion is that, no matter how tightly packed the DRG, it is impossible to create enough contacts.

As a result, our findings suggest that none of the leading hypotheses explain the phenomenon.

## Introduction

Neurons of dorsal root ganglia (DRG) are the first-order sensory neurons, tightly packed into clusters positioned bilaterally along both sides of the spinal cord. These clusters are practically devoid of synapses, and the neurons themselves have a distinctive morphology: each has only two outgrowths entering the soma side by side, creating the characteristic pseudo-unipolar form of these cells [1]. Beyond neurons, DRGs are composed of satellite glial cells (SGCs) that play roles similar to astrocytes in the central nervous system [1], [2]. SGCs regulate the microenvironment of DRG neurons and guide their neurogenesis toward the pseudounipolar form [3]. These glia cells create capsule-like structures around each neuron, effectively isolating it from direct contact with neighbors [2], [4]. For simplicity, we refer to a neuron and its surrounding glial cells as a module. SGCs surrounding a particular neuron sometimes connect to one another via gap junctions [5], while glia cells of distinct capsules typically separated by connective tissue [1]. In pathological conditions or with age, however, this isolation may break down, with SGCs forming syncytium-like interconnected networks across multiple modules [5], [6], [7]. The exact triggers for this alteration remain unclear, though it appears to contribute to neuropathic pain pathologies in some cases [8].

According to the classical view [1], DRG neurons do not perform complex neural computations, merely summating signals from each neuron’s dendritic tree. In this traditional perspective, DRG neurons function as single-channel pre-amplifiers that filter incoming ‘raw’ signals and transmit them further only if they exceed specific thresholds. However, this perspective has been challenged by discoveries in the last two decades. Particularly a phenomenon of ‘cross-excitation’ was observed, during which nearby neurons demonstrate significantly increased excitability when their neighbors are stimulated [9], [10]. perhaps, this interaction results in some sort of convolution-like process, allowing for better preprocessing of sensory stimuli. This conclusion, if confirmed, will mean that processing of sensory information happens even before the signals enter CNS.

Another significant characteristic of this phenomenon is its temporal profile: the increase in excitability does not occur instantaneously (as would be expected from ephaptic effects mediated by electromagnetic fields) but within a time window of 0.5 to 1 second after the initial stimulus [10]. This delay suggests the involvement of diffusion-mediated transmission mechanisms, assuming sufficient propagation speeds for the mediators.

Notably, the distance between glial cells and neurons within a module is remarkably small, approximately 20 nm [2]. Since membrane potential heavily depends on concentration differences between intracellular and extracellular spaces [11], this minimal volume surrounding the cell underscores the critical importance of SGCs in forming both resting and action potentials.

Despite the significance of cross-excitation, the precise mechanisms underlying this phenomenon remain unclear, with no single theory receiving decisive supporting evidence. One leading hypothesis suggests mediation through gap-junctions of nearby modules. The overall scheme looks as follows: when a neuron of the one module fires, it releases Ca or ATP into its surrounding space, and these substances are picked up by SGC. If SGC has connections to another module’s SGC, it will transmit either these substances or signaling molecules (P2X3, in case of ATP) to them, causing changes in SGC-sheath of a nearby neuron. Though this hypothesis was successfully verified in numerical models [12], [13], the number of gap-junctions between different neurons’ glia is stated to be very low (from 0 and up to 6.6% occurrence, [5], [6]). Although this occurrence was shown to be orders of magnitude higher in pathological condition [5], [6], the cross-excitation phenomena was initially discovered in intact dorsal ganglion. This raises the question, is this amount of gap-junctions enough to mediate the effect in the normal state?

Experimental work has demonstrated significant increase 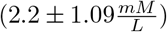 in extracellular K+ levels in tonically stimulated dorsal root ganglia [10]. These changes may result from potassium diffusion either through the extracellular space or within glial cells. The possibility of the former was experimentally shown, as ions can indeed pass from one neuro-glial cleft to another [14]. In both cases, such changes may be responsible for nearby neurons’ excitation changes, as potassium is crucial for action-potential formation. Utzschneider’s experiments included measuring potassium concentrations around a neuron as well as its membrane potential. Changes in membrane potential of the amplitude of 3.6 3.5 mV accompanies those concentration shifts. It was not estimated whether these ion changes alone were the reason for potential change or vice versa. Additionally, colleagues [10] suggested that the effect is of populational nature - not one but several nearby neurons firing together are able to shift excitability of the target one.

An alternative idea proposes a chemical mediator-based effect, as changes in ion concentration alone might be insufficient, but even small amounts of signaling molecules could trigger the observed effects. It was extensively researched during the last decade, resulting, among others, in the hypothesis of ‘sandwich-synapse’ [15], [16]. This idea, although verified by clear molecular sightings, explains the cross-excitation phenomenon in a very particular configuration of neurons - when two somas are located on both sides of one SGC (hence the sandwich name). Occurrence of such a situation is stated to be around 6-20% of all DRG neurons, leaning to the left side of this interval, as stated by the authors themselves [15]. It raises the question whether this chance is enough for 87-95% of neurons to be interconnected.

Lastly, it can be hypothesized that the cross-excitation phenomenon can be attributed to some complex electro-magnetic (ephaptic) interaction between DRG neurons. Although such interactions indeed were discovered in the peripheral nervous system [17, 18] and are enhanced by limited space (for example, tightly packed nerve [18]), they are still insufficient to cause significant depolarization [17]. As for electromagnetic interactions betwen neuron bodies, this interaction are extremely small even in compartment as tight as human cortex [19]. Consider that, unlike DRG neurons, cortical neurons are not isolated from each other by glial sheaths [20], further decreasing the possibility of such interplay.

Additionally, a problem of mechanisms’ interaction shall be mentioned. Of course, it is a usual way of thinking, that one effect is explained by one mechanism; in the end, it is a natural following from Occam’s razor. However, it is often not the case with living systems, especially as complex as a central nervous system of mammals. It is certainly possible that the cross-excitation effect may be mediated via a combination of mechanisms. We touch this idea in the ‘Geometrical simulation’ part, in which we analyze the CE connection from probabilistic perspective. However, to fully estimate it, a separate research shall be conducted, as clear and sound arguments shall be given for two different mechanisms explaining one phenomenon.

## Methods and Models

All described simulation were performed using Python programming language and the following libraries: SciPy [21], NumPy [22], Numba [23], Skfem [24], SkOpt [25], Pandas [26].

We aimed to computationally verify both diffusion-based and gap-junction-based hypotheses. For the diffusion hypothesis, we tried to recreate the system as fully as possible, allowing for diffusion of K+ between two modules. As for gap-junction hypothesis, we used data on their occurrence in DRG as well as Monte-Carlo sampling to analyze statistical plausibility of the interaction.

### Diffusion simulation

In recent years, a comprehensive model of DRG neuron was developed [27] - we recreated it 1 to 1 and combined it with a model of satellite glial cell [12] to recreate a complete model of DRG module. This model is an expansion of the classical Hodgkin-Huxley model [28], with all currently discovered channels and pumps of DRG neurons included. We expanded it, by allowing for dynamic ion changes by converting each ion’s flux into concentration changes according to formula 1.

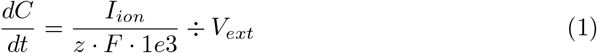

where 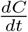-concentration changes, in 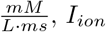-electric current, carried by specific ion through the whole surface of cell, in mA, z - unitary charge of the ion, F – Faraday constant 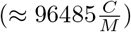, *V*_*ext*_ - volume of extracellular space, in L. To fully assess all the ions’ effects, channels which are traditionally considered ion non-specific (i.e., leakage channels, TRPM8 channels, etc) were made specific, using standard permeability relations from [11, 29, 30]. Volumes were calculated as nested spheres - a neuron, with radius 20 micrometers, nested inside a spherical SGC sheath with thickness of 3 micrometers. Neuron was separated from a sheath with a cleft of 20 nanometers, in which ion concentrations were updated dynamically by both glia and neuron.

The resulting multicellular model produced dynamics similar to that of a real DRG neuron, when put in normal physiological conditions (T = 37C, ion concentrations similar to those in [27]). Example of action potential generated by the model in response to adequate stimulus can be seen at 1, while potassium concentration changes depicted on 2. 3, 4 depict responses to pulse-train stimulation.

**Fig 1.**
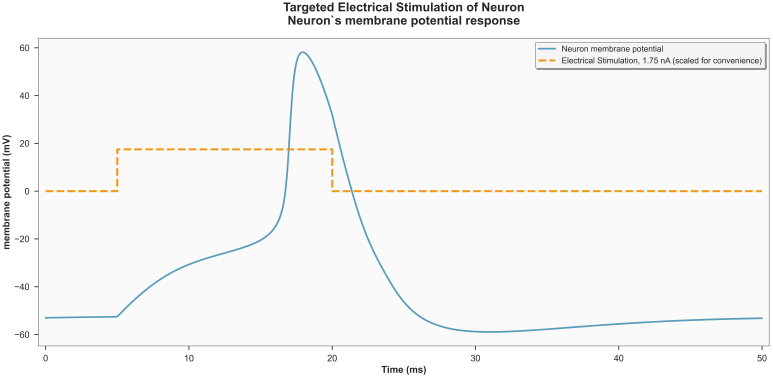
Singular action potential of simulated DRG neuron.

**Fig 2.**
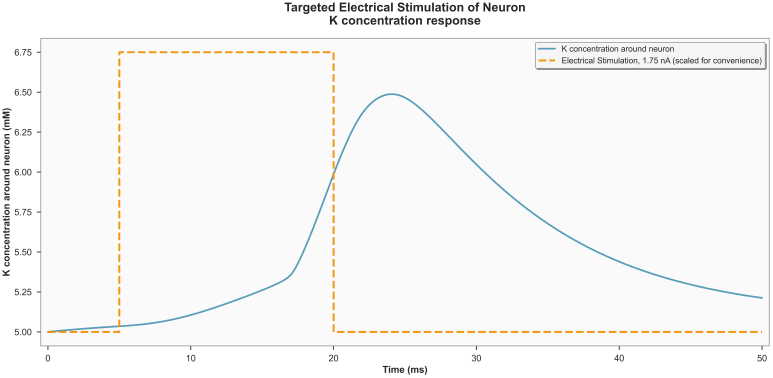
Changes in K concentration in cleft between neuron and SGC during singular stimulation.

**Fig 3.**
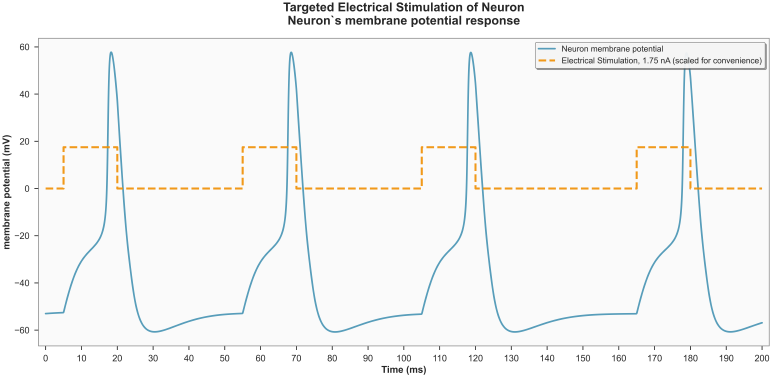
Multiple action potentials of simulated DRG neuron.

**Fig 4.**
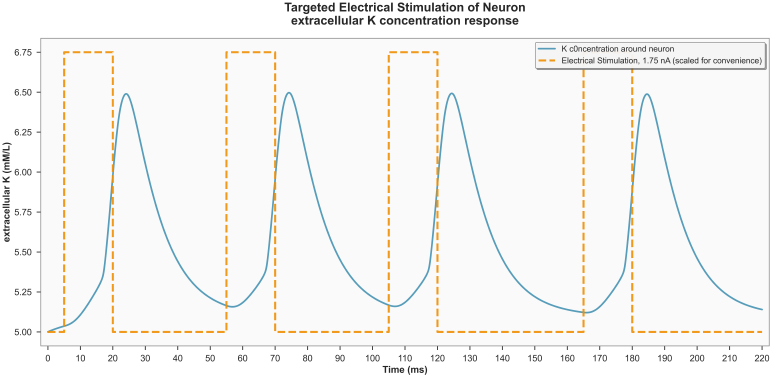
Changes in K concentration in cleft between neuron and SGC during singular stimulation.

#### 0.0.1 Experiment 1

The first step to test the diffusion paradigm was to assess how much the membrane potential is changed by changing K concentration around a neuron. Experimental data suggest a change of 3.6 ± 3.5 mV membrane potential and of 2.2 ± 1.09 mM *K*_*o*_ increase when the neuron is affected by cross-excitation. There is a reasonable approximation for membrane potential - a Goldman-Hodgkin-Katz equation [11]:

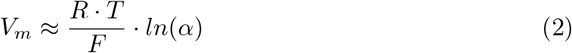

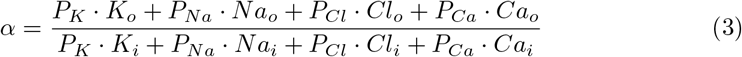

Where R - universal gas constant 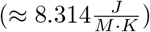, T - temperature (taken as 293.15 K, in-vitro temperature), F - Faraday constant 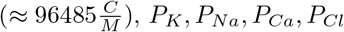, – relative permissibilities of respectable ions, *K*_*o*_, *Na*_*o*_, *Cl*_*o*_, *Ca*_*o*_ - concentrations of ions outside the membrane, in mM, *K*_*i*_, *Na*_*i*_, *Cl*_*i*_,*Ca*_*i*_ - concentrations of ions inside the cell, in mM. Substituting concentrations from [27] and taking standard values for relative permissibilities from [11], [30], we get:

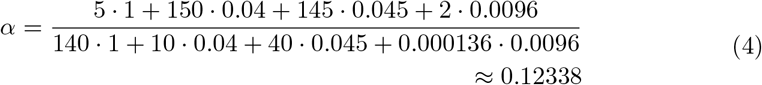

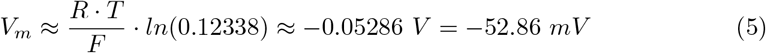

Thus, in intact condition, *V*_*m*_ is expected to be around −52.86 mV, which is a reasonable approximation (the model produces values of −53 mV). For experimentally observed shifts, this GHK estimation becomes −49.49 mV, thus the effect of increased potassium concentration is +3.37 mV. Therefore, the change in membrane potentials aligns with K concentration changes, if approximated with Goldman-Hodgkin-Katz equation.

Therefore, we estimated the membrane potential shift in a comprehensive DRG neuron model with fixed ion concentrations to assess how much the potential would increase if neither SGC, nor a neuron attempted to stabilize it. In this experiment, we compared the membrane potential of a DRG neuron under normal conditions (*K*_*o*_ = 3.6 mM) against a neuron with increased *K*_*o*_ (up to 5.8 mM). Throughout the simulation, ion concentrations remained fixed, and we allowed the system to adapt to the change. The resulting *V*_*m*_ of a neuron in increased *K*_*o*_ after 15 ms stabilized around −50.48 mV (compared to −53 mV under intact conditions). Third, we analyzed how the system behaves when concentrations are allowed to change dynamically. In this scenario, all concentrations are calculated dynamically and are affected by both neuron and glia, as well as diffusion into connective tissue. Thus, only the initial concentrations were set to increased values. To compare the dynamics, we subtracted the time series of *V*_*m*_ after the potassium concentration change from the series before the change, obtaining a difference graph. The maximum observed difference was 1 mV, and on average, the membrane potential increased by 0.746 ± 0.239 mV. In actual experiments, scientists observed neurons with different resting state potentials (*−*60 ± 7.4 mV, not higher than *−*50 mV) [10]. To recreate this situation, we varied the passive leakage channels’ electrochemical potential (*E*_*pas*_) to set different resting state potentials (RSP) for different neurons, exploiting the direct connection between these parameters in the Hodgkin-Huxley formalism. We sampled 100 neurons with membrane potentials of *−* 61.36 ± 6.05 mV and analyzed how their *V*_*m*_ would change after increasing *K*_*o*_ (the increase was again sampled from real data and was 2.2 ± 1.09 mM). Membrane potentials of neurons before and after changes were compared, revealing an increase of 0.738 ± 0.292 mV. Thus, the results are quite controversial: at first glance and in simplest approximation, the changes are sufficient; however, the more biologically plausible the situation we analyze, the bigger the difference between experimental and theoretical data becomes.

#### 0.0.2 Experiment 2

We then decided to recreate the full scheme of cross-excitation. By connecting two neuron modules with diffusion space, we can estimate the amplitude of concentration changes produced by one firing neighbor. Even if the effect results from the summation of activity from several neighbors, the case with two neurons should show proportionally smaller changes.

We performed the simulation according to the following scheme: two neuron modules, each containing a neuron and nearby glia. The extracellular space of each neuron (internal space of a module) is connected to a 1D diffusion space that imitates connective tissue between modules. At each simulation step, K+ concentration is first updated by the neuron and its glia, then diffused within this space.

Diffusion was calculated according to simple Fick’s law:

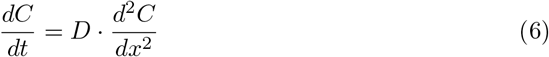

where C - concentration of an ion, x - distance in space, cm, D - diffusion coefficient, in 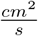 (*D* = 5 *·* 10^*−*9^, derived from [31]). During 20 seconds of the simulation, one of the neurons is stimulated with 20 Hz 10 ms long square pulses of 1.75 A amplitude to imitate the experimental conditions. We compare the changes in membrane potential of a second, not-stimulated neuron, to the case in which no stimulation is applied to both cells. The resulting difference is in order of 5 *·* 10^*−*4^ 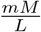or of 6 *·* 10^*−*4^ *mV* (figs. 5, 6).

**Fig 5.**
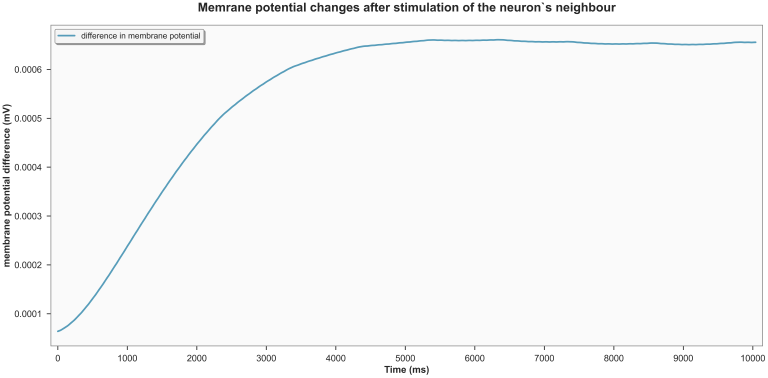
Changes in membrane potential when a neighboring neuron is stimulated.

**Fig 6.**
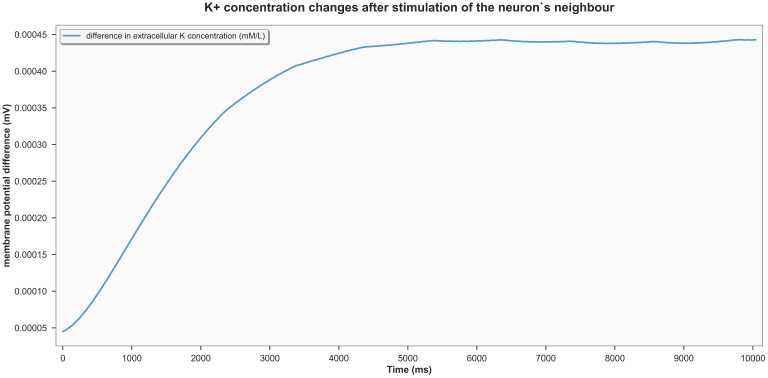
Changes in K+ concentration in neuron-SGC cleft when a neighboring neuron is stimulated.

**Fig 7.**
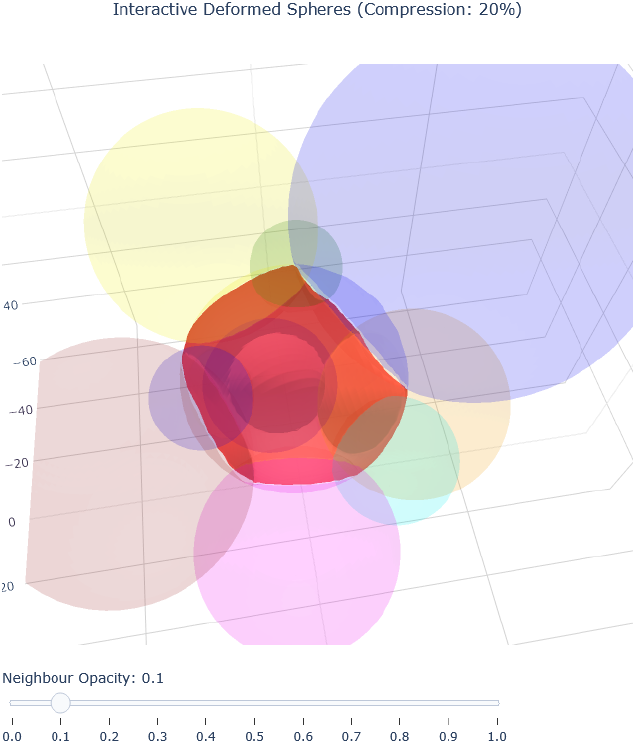
Example of generated spheres’ positioning in 3D space

**Fig 8.**
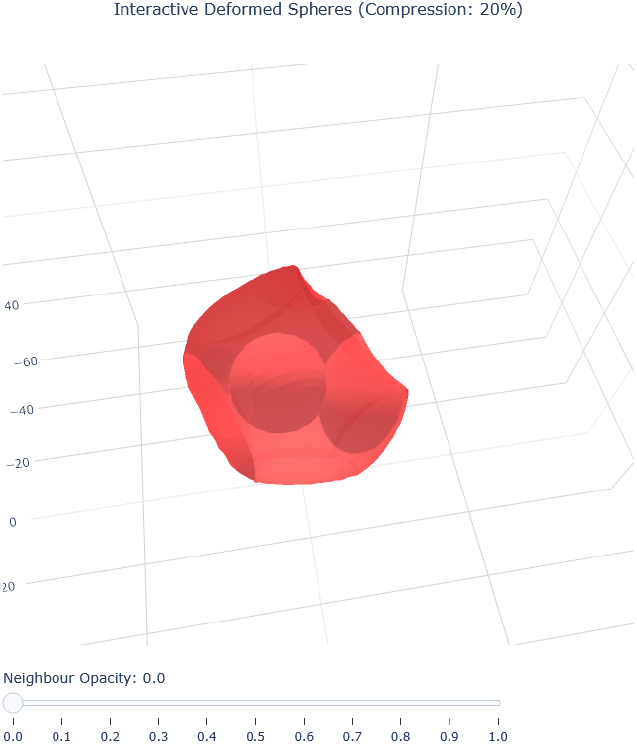
Example of generated spheres’ positioning in 3D space (with opaque neighbors)

**Fig 9.**
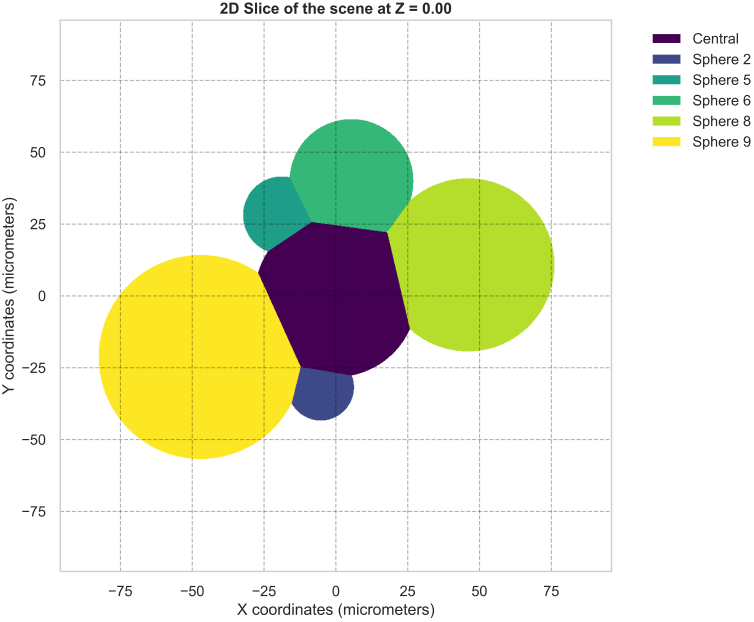
Example of 2D slice through generated scene

**Fig 10.**
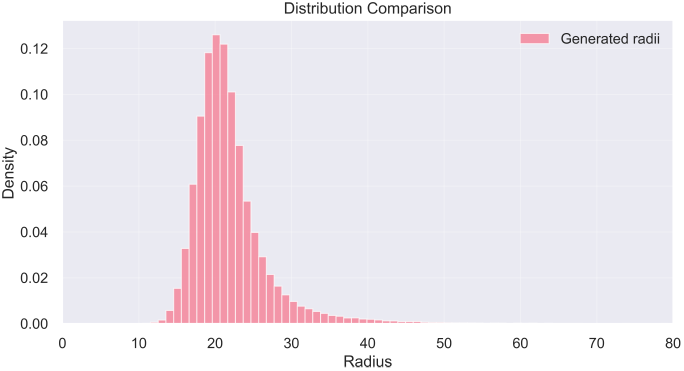
Distribution of generated spheres’ radii

**Fig 11.**
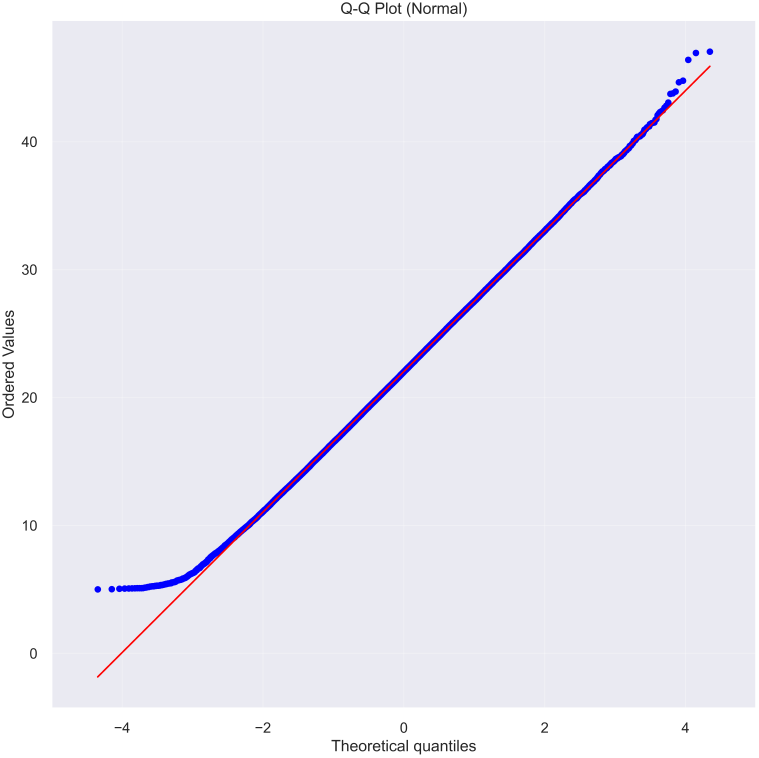
Quartile-Quartile plot for generated spheres

**Fig 12.**
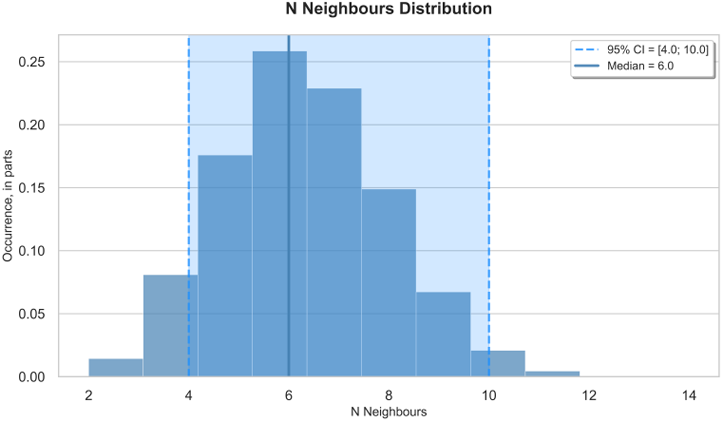
Quartile-Quartile plot for generated spheres

**Fig 13.**
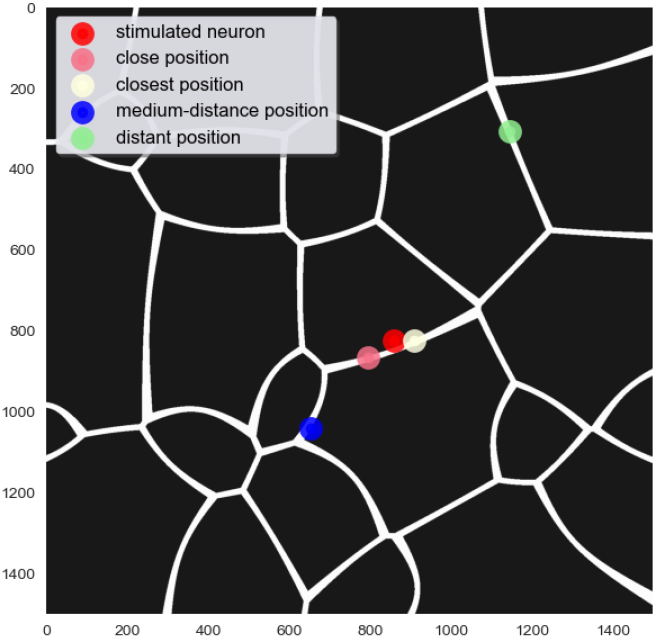
An example of simulated labyrinth of extracellular space

**Fig 14.**
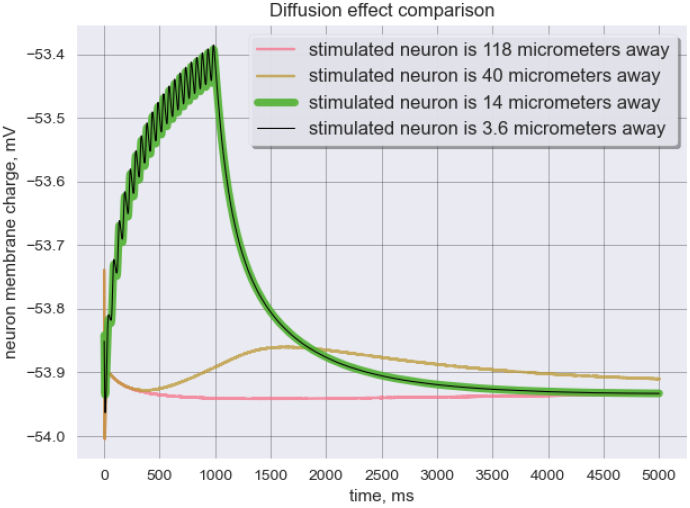
Graph of membrane potential of the affected neuron

**Fig 15.**
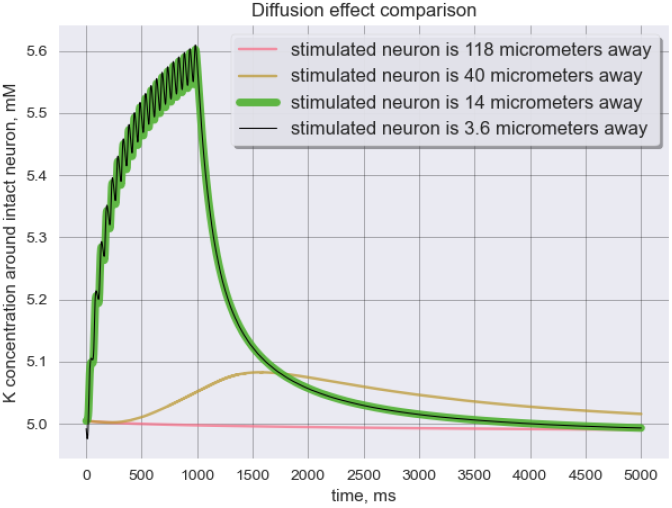
Graph of K concentration changes around the affected neuron

**Fig 16.**
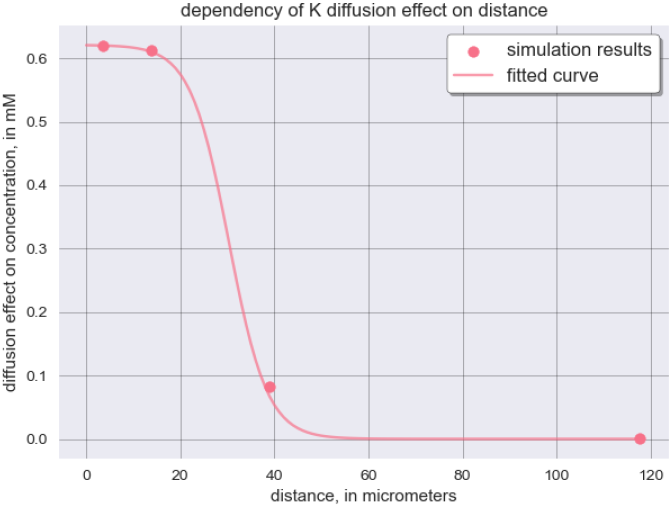
Graph of how diffusion-based interaction depend on distance

**Fig 17.**
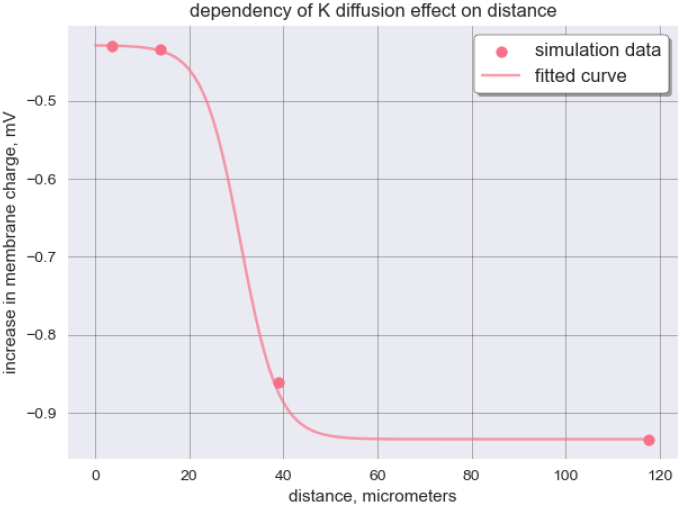
Graph of how diffusion-based interaction depend on distance

This is many orders of magnitude lower than necessary for any significant excitability change - even if the effect of one neuron is 1 · 10^4^ times weaker than the groups’ one, it is still not enough to produce an observed amplitude of cross-excitation.

However, we also ran the model with two simultaneously stimulated neighbors (as this is the maximum amount of immediate neighbors in one-dimensional space space). Thus, we roughly estimated how the effect of K concentration change scales with the neighbors’ number. For one dime case, it seems to be linearly dependent, as 2 stimulate neighbors produced exactly twice the change created of a one neighbor. Thus, to reach the observed levels, the number of direct neighbors shall be of the order of thousands (*≈* 3 · 10^3^, assuming linear scaling), which, as will be shown later, is impossible. We will discuss further the problem of the number of immediate neighbors in the next chapter. One of the important parameters is the distance between neurons in diffusion space.

Note that it is not a shortest distance between two modules’ sheaths - it is a way between openings of those sheaths which allow for coupling through diffusion. We estimated the model on different distances between modules - and found no significant difference in Vm changes in biologically possible range of distances (from 0.005 cm to 0.0005 cm, differences between ranges *<* 1 · 10^*−*16^ mV, that is, in the range of numerical error). Thus, we turned to other hypotheses in search of the explanation of the cross-excitation phenomenon.

### 0.1 Geometrical simulation

Next, we decided to test the gap-junction mediated hypothesis. As it was already shown that if two modules are connected by gap-junctions, cross-excitation is possible. The occurrence of GJ connecting different neurons’ glial sheath is known and stated to be from 0 to 6.6% in norm [5, 6, 32].This number means the probability of two modules being connected. On the other hand, the occurrence of cross-excitation of 95% means that, if we pick one neuron, there is a 95% probability of it being affected by neighbors. In case of the gap-junction hypothesis, this becomes a discrete value - there should be at least one connected neighbor in 95% of cases. Thus, we can state that, if the occurrence of cross-excitation is 95% percent, there is exactly 95% percent chance that, if we pick a random neuron, there will be a connected neighbor. Note: it is not the same as the chance of a particular link (particular GJ connection) existing, it is a chance for a particular neuron to be coupled to one or more of its neighbors.

Now, the question is: knowing the probability of two neighbors to connect, how many neighbors is necessary for us to observe the stated occurrence of cross-excitation.

For now, taking only publications where such connections were observed in norm, we assume a 4.65% chance of them being connected (probability of particular connection to exist).Probability of them not being connected = 95.35% (probability for particular connection to be absent, call it *p*_*a*_). Therefore, if we pick a module and then search through surrounding modules for a connected one, probability is defined as:

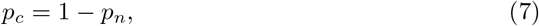

where *p*_*c*_ - probability to find at least one couple, *p*_*n*_ - probability not to find a couple, which is equal to:

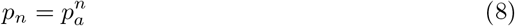

where n - number of nearby modules, since the only case when we do not find any coupled neighbors is the case when all connections happen to be absent. Now we can directly estimate how many neighbors are required for cross-excitation occurrence of 95%:

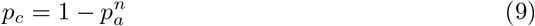

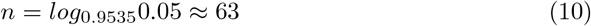

And for cross-excitation occurrence of 87%:

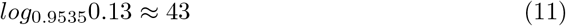

Here we make another assumption, since we are concerned about populational properties and not single cells, we can approximate a neuron as a sphere. The task becomes kissing sphere problem - yet an unsolved mathematical problem about maximum number of neighbors that is possible to fit around a sphere. For equally sized spheres in 3 dimensions the solution is 12 [33], so the chance of cross-excitation appearing between a neuron and any of its neighbors is:

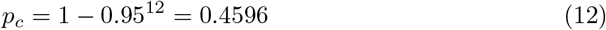

But in our case spheres vary in size - there is a distribution of neuron radii, with mean of 24.5 micrometers and standard deviation of 5 [34]. Firstly, we shall estimate the upper-bound case - when the central neuron is huge and all of its neighbors are small. There is an upper limit approximation for such a case [35]:

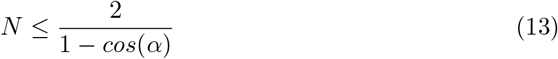

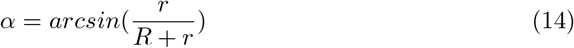

Where N - maximum possible number of neighbors, r - radius of neighbors (smaller than central), R - radius of a central sphere. Taking *R* = *mean* + 3 ·*SD* = 39.5 and *r* = 5 we can estimate N neighbors:

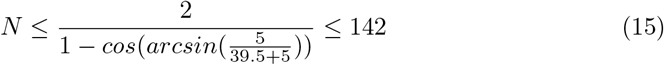

Thus, there is a mathematical possibility that we can indeed observe the necessary number of neighbors, if we happen to pick a very large neuron surrounded by very small ones. Now, the question becomes: what is the probability for us to get such a structure? We shall estimate how probable it is to get such a boundary case, i.e. to estimate the distribution of neighbors. We can do this by performing Monte-Carlo simulation [36] - sampling neighbors with the radius taken from the known distribution until all the vacant places around one neuron are taken. Repeating this enough times will give us an estimation of N neighbors distribution. The outline of the algorithm:

1. create a central node with the radius sampled from real normal distribution
2. create a neighbor with radius sampled from real normal distribution
3. find a random direction and place it nearby
4. check, whether it overlaps with already existing neighbors
5. if yes, go to 3, if no - go to 2
6. repeat for set number of cycles for set number of simulations

However, this approach shifts the distribution of neighbors’ radii - the more modules are placed, the more probable it is to fit in smaller modules. We can avoid this by limiting the number of tries, after which the script stops trying to fit in a particular neighbor of this module and goes to the next one. This approach still shifts the distribution and, more importantly, changes its shape. Thus, we introduced an inflation coefficient, a special coefficient that increases the radius of a sphere we are trying to fit depending on the number of consecutive unsuccessful iterations. We fitted this parameter using Bayesian fitting [37], as it is very suited for random-process’ parameters fitting.

We ran the simulation for 1 ·10^5^ times, generating this number of ‘central’ neurons and for each of them trying to fit each of the drawn neighbors for 50 times with inflation coefficient of 0.02, until considering all the vacations taken. Further, if we ‘press-in’ neighboring spheres into the central one - imitating tight packaging - the solution resembles the situation observed in real DRGs 7 (8 demonstrates results with completely transparent neighbors). 9 demonstrates a 2D slice of this exact situation along a horizontal plane and compares it with a real slice. This is purely illustrative touch to show that, not only statistical and linear metrics of real DRG are recreated, but, without altering those metrics, we may recreate semi-natural view of ‘slice’, observed in histology.

The distribution of radii we observed (10, 11) was not statistically different from the real distribution (KS-test, p-value¿0.05, 24.55 against generated 24.027 5.02).

As for the immediate neighbors, their number necessary to allow 95% connections (63) is far beyond 95% percentile (which is 10), as well as 99.9% percentile (11), thus this situation is statistically impossible (12).

We also can estimate other proportions - in other papers [10] the number of 87% connected neurons is found. This probability of connection would require 43 immediate neighbors. The value is still beyond 99% percentile. There were also several articles proving that if two neurons lay on two sides of one SGC, or enveloped by one SGC, there is a chemical signaling possible [15, 16, 38]. It was estimated that around 10% of neurons are found in such conformations. Thus, this chemical mechanism to be present in 95% of observations requires 29 neighbors on average - still an impossible situation. Taking the upper margin, despite researchers claiming it shall be closer to the lower boundary, gives us 15% of connections - approximately 19 neighbors required. If we assume that both mechanisms are present - that at least one type of connection should be present for CE to work, we can give the probability of 0.0465 + 0.1 = 0.1465, for which we will still need 18 neighbors.

if we take the maximum mentioned numbers - 0.066 for gap-junction occurrence, 15% for chemical ‘sandwich-synapse’, we get 21.6% neurons connected, requiring only 14 neighbors. It is still beyond the 99% percentile. Therefore, even if this situation is theoretically possible, it is a highly unlikely - less than 5% of neurons would be connected by cross excitation, meaning that out of 18 modules [9] had tested, only one would show the cross excitation (meaning that only one would have had enough neighbors to find a connected one).

### 0.2 Combination of diffusion and geometrical simulation

In the diffusion simulation part, we simulated the process of potassium diffusion between two neurons. However, in a real dorsal root ganglion, the extracellular space is not devoid of obstacles; quite the opposite - due to the very dense packing of neuronal modules, the extracellular space suitable for diffusion resembles a complex labyrinth rather than an open volume.

Using weighted Voronoi diagrams [39] and the approach developed in the second step, we simulated DRG space filled with neuronal modules with a realistic distribution of radii. We then moved the modules apart to create an inter-modular distance of 7.5 micrometers. This value was chosen as an approximation based on anatomical considerations: while no direct measurements of inter-modular spacing have been reported in the literature, we can reasonably speculate that, since this space must accommodate connective tissue and the smallest blood vessels (capillaries), it should be at least 5 to 10 micrometers on average [40]. Based on this reasoning, we generated a labyrinthine geometry with a distance of 7.5 micrometers between walls. Since we are modeling a widespread phenomenon, local architectural variations are of limited importance - at a sufficiently large scale, the system will converge to average parameters, and these average parameters must be conducive to cross-excitation; otherwise, the phenomenon would not be as prevalent as observed experimentally.

We repeated the experiment from the first step: simulating one neuron being stimulated with an action potential-generating current while its direct neighbor remained at rest. The neurons were positioned as immediate neighbors, with each connected to the extracellular labyrinth through a gap in the glial (satellite cell) envelope. The dimensions and characteristics of these gaps are not well-documented in the literature; it is simply stated that “…some portions of the neural surface seem to be without a satellite cell covering…” [2]. Therefore, we systematically analyzed the relationship between gap size and the magnitude of cross-excitation via potassium diffusion. Varying the gap size from 5 to 20 micrometers in width, we observed exactly zero difference in the effect amplitude. This is likely due to the fact that, when time is sufficiently big, ionic flux through surface becomes close to zero, no matter section through which the flux happens.

It remains unknown whether there is one or several such gaps per neuron, but this distinction is non-essential from a modeling perspective, as multiple smaller gaps are functionally equivalent to a single larger gap with the same total area. However, the spatial positioning of the gaps is critical: if the gap of the stimulated neuron faces the gap of the intact neuron, the cross-excitation effect will be substantially stronger than in other spatial configurations. Consequently, we also analyzed the dependency of effect amplitude on gap positioning, specifically the distance between gaps measured as the shortest path through the labyrinthine extracellular space (see 14, 15, 16, 17). Distance was calculated as an optimal path inside the labyrinth, using Breadth First Search algorithm [41]. The resulting amplitude of concentration change did not exceed 0.5 mM, when two modules’ gaps are opened practically into each-other. This change corresponded to shift in RSP of *≈* 0.4 mV.

Our results demonstrate that none of the tested geometric configurations produce cross-excitation effects of sufficient magnitude to explain experimental observations—even in the most favorable case where gaps directly face each other. The effect reaches physiologically relevant amplitudes only when the two neuronal modules’ interior spaces are located so close they can be consiered practically connected. This situation is analogous to the case where two neurons share a common glial envelope. Unfortunately, such configurations are extremely rare (around 10% of cases [15]. Thus, we conclude that even when the volume of the diffusion-mediating extracellular space is reduced by approximately an order of magnitude (as in our model), the resulting cross-excitation effect remains insufficient to explain experimental observations.

## Results

### 0.3 The Diffusion Hypothesis is Insufficient to Explain Cross-Excitation

#### 0.3.1 Local Potassium Increase Fails to Cause Sufficient Depolarization

To assess whether localized potassium accumulation could mediate cross-excitation, we simulated extracellular K+ increases from physiological levels (3.6 mM) to pathologically elevated concentrations (5.8 mM) in isolated neuron modules. The Goldman-Hodgkin-Katz approximation predicted a membrane potential shift of approximately 3.2 mV under these conditions. Further, when implemented in our comprehensive dynamic model accounting for neuronal and glial ion regulation, the actual depolarization was substantially smaller at 0.75 ± 0.29 mV (n=100 neurons with varied resting potentials of *−* 61.36 ± 6.05 mV). This represents less than 10% of the experimentally observed 10 mV depolarization characteristic of cross-excitation events, indicating that local K+ accumulation alone cannot account for the phenomenon.

#### 0.3.2 Diffusion Between Modules is Negligible

We next examined whether ionic diffusion between spatially separated neurons could mediate cross-excitation by simulating K+ release from an active neuron and measuring the resulting potential change in an adjacent, unstimulated neuron. Our two-module simulation, incorporating realistic diffusion dynamics through connective tissue, revealed membrane potential changes on the order of 10^*−*8^ mV in the receiving neuron. This negligible effect demonstrates that passive ionic diffusion between discrete neuronal compartments is functionally irrelevant for cross-excitation, even when accounting for the cumulative effects of multiple firing neighbors.

Moreover, even if diffusion is simulated in tightly-packed space with realistically shaped neurons separated by extracellular space as tight as 7.5 micrometers, diffusion-mediated effect is observable (in orders of hundredth to tenth of mM) only in extreme proximity. Such proximity can be obtained if two somas are either located in one envelop or each of them have a gap in the envelope opened towards the neighbor. This conclusion is of extreme importance: it heavily limits the number of neighbors which are significantly affecting the target neuron. This, coupled by median number of immediate neighbors being 7, makes the diffusion-based transmission impossible.

### 0.4 Structural Connectivity Hypotheses are Statistically Implausible

#### 0.4.1 The Required Number of Neighbors Exceeds Physical Possibility

To evaluate structural connectivity hypotheses, we calculated the number of neuronal neighbors required to achieve the observed 87-95% prevalence of cross-excitation through known coupling mechanisms. Given the reported 4.65% probability of gap junction formation between adjacent neurons, achieving 95% cross-excitation probability would require each neuron to have approximately 63 neighbors, while even the lower observed prevalence of 87% would demand 43 neighbors per neuron. We tested these requirements against anatomical constraints using Monte Carlo simulations of DRG neuron packing based on realistic cell size distributions and tissue geometry. Our analysis revealed a median of only 7 neighbors per neuron, with 95% of neurons having 13 or fewer direct neighbors. This represents a fundamental mismatch between the structural requirements of gap junction-mediated cross-excitation and the physical reality of DRG organization.

### 0.5 Combining Known Mechanisms Does Not Resolve the Discrepancy

We further evaluated whether combining gap junctions with “sandwich synapses” (ephaptic coupling through shared satellite glial cell ensheathment) could reconcile this discrepancy. The combined probability of either mechanism occurring between neighboring neurons is approximately 14.7%. However, even with this increased coupling probability, achieving 95% cross-excitation would still require approximately 18 neighbors per neuron. This number exceeds the 95th percentile of our packing simulations, indicating that known structural mechanisms, either individually or in combination, cannot explain the high prevalence of cross-excitation observed in healthy DRG tissue. Together, these findings demonstrate that neither diffusion-based nor chemically mediated mechanisms can explain the robust cross-excitation observed in the dorsal root ganglia, suggesting the involvement of alternative, yet unidentified coupling pathways.

## Discussion

According to our results, neither the diffusion-based nor gap-junction-mediated hypotheses of the cross-excitation mechanism are statistically and/or biophysically possible. The diffusion mechanism produces a negligible effect that cannot explain the observed fluctuation of extracellular K+ and, consequently, the neuron’s membrane potential. Furthermore, even when diffusion is calculated through realistically crumpled space - where the actual available volume is more than 10 times smaller - the effect remains insignificant.

Simultaneously, gap-junction-based cross-excitation is impossible due to the extremely low probability of such connections between neurons. For this probability to support the observable effect amplitude, each neuron would require a geometrically impossible number of immediate neighbors. While our results indeed contradict the explanations initially proposed by Utzinger et al. and Devor et al., they should be seen as a direct extension of that research. The original hypotheses were logical and clear interpretations of freshly acquired evidence at the time. For instance, the diffusion hypothesis was a clear and direct conclusion from the observed increase of extracellular potassium following neighboring neuron firing. It was based upon a robust fundamental understanding of the neuron firing mechanism. However, this observation merely tells us that the concentration increase happens, not the mechanism behind it. Considering the extremely complex homeostatic neuroglial interactions, it becomes impossible to derive the answer solely from the fact of the potassium increase. By isolating the variable, we demonstrate that the resulting depolarization from diffusion alone is several orders of magnitude smaller than required, suggesting the observed K+ flux is likely an epiphenomenon rather than the cause.

Similarly, the gap-junction hypothesis was grounded in strong evidence of existing physical connections between neuron modules. However, the critical question of the prevalence of such connections is difficult to resolve anatomically, since one cannot directly estimate whether the number of connections is “enough” or “not enough” simply by analyzing histological data. We addressed this lacuna, once again extending what was already discovered experimentally.

Of course, the present work is heavily limited in terms of how many hypotheses we could test in a purely theoretical setting. With it, we merely hoped to outline an intriguing inconsistency found by others and, perhaps, to draw the attention of researchers to this phenomenon. As with any modeling, we make numerous assumptions, some more speculative than others. For example, the approximation of neuron shape to a sphere is widely used and can be easily attributed to the average form of the cell being roughly spherical. However, for the size of the extracellular space between DRG neurons, we could know neither its actual width nor shape. It is possible that the volume available for diffusion differs from what we stated, but not significantly: ultimately, we are working with widespread phenomena, and thus averaging and estimation based on mean values is applicable. Otherwise, if the effect required, for example, specific ganglia geometries, the effect would either be rare or such conformations would be widespread.

All in all, we strongly believe that deep understanding of underlying mechanisms of cross-excitation would bear both fundamental and practical outcomes.

From a fundamental physiological perspective, understanding the exact mechanisms of DRG neuron interactions could provide new insights into how sensory information is processed at the most basic levels. If the interaction is so common, each impulse from a DRG neuron inevitably carries information from its neighbors. What is the proportion of this cross-transmitted information? How does the nervous system utilize this additional information? We believe that answering these questions offers a new perspective on the principles of sensory perception. It would be an intriguing endeavor to model this process, as such prevalent and strong interaction should greatly alter the sensory information transmitted via DRG. In our future work, we plan to dive deeper into this, examining what exactly is altered, how it can be described, and what computational benefits this particularity brings for the nervous system.

From a practical point of view, our limited understanding of cross-excitation prevents us from fully estimating its interaction with spinal electrical stimulation techniques, one of the most promising approaches for rehabilitation after spinal cord injury. As spinal cord stimulation (both invasive and non-invasive) is applied to DRG as well - since dorsal roots also happen to be within the stimulated radius - the processes occurring inside the ganglion become important. If the occurrence of cross-excitation is indeed this high, it is inevitably involved in neurostimulation. It has already been hypothesized that, in terms of neurogenic pain treatment, stimulation of DRG instead of the cord itself is promising [42, 43]. Speaking generally, these spinal stimulation techniques have already shown remarkable results in helping patients recover from severe traumas [44, 45]. However, it may be possible to achieve even better rehabilitation by adjusting for antidromic effects and their interplay with cross-excitation [46]. It is also possible that such knowledge will enable clinically applicable direct stimulation of DRG [47].

## Conclusion

## Supporting information

**S1 Fig. Bold the title sentence**. Add descriptive text after the title of the item (optional).

**S2 Fig. Lorem ipsum**. Maecenas convallis mauris sit amet sem ultrices gravida. Etiam eget sapien nibh. Sed ac ipsum eget enim egestas ullamcorper nec euismod ligula. Curabitur fringilla pulvinar lectus consectetur pellentesque.

**S1 File. Lorem ipsum**. Maecenas convallis mauris sit amet sem ultrices gravida. Etiam eget sapien nibh. Sed ac ipsum eget enim egestas ullamcorper nec euismod ligula. Curabitur fringilla pulvinar lectus consectetur pellentesque.

**S1 Video. Lorem ipsum**. Maecenas convallis mauris sit amet sem ultrices gravida. Etiam eget sapien nibh. Sed ac ipsum eget enim egestas ullamcorper nec euismod ligula. Curabitur fringilla pulvinar lectus consectetur pellentesque.

**S1 Appendix. Lorem ipsum**. Maecenas convallis mauris sit amet sem ultrices gravida. Etiam eget sapien nibh. Sed ac ipsum eget enim egestas ullamcorper nec euismod ligula. Curabitur fringilla pulvinar lectus consectetur pellentesque.

**S1 Table. Lorem ipsum**. Maecenas convallis mauris sit amet sem ultrices gravida. Etiam eget sapien nibh. Sed ac ipsum eget enim egestas ullamcorper nec euismod ligula. Curabitur fringilla pulvinar lectus consectetur pellentesque.

## Acknowledgments

This work was supported by Saint Petersburg State University (projects No.125022102790-5 to P.M.), and by the State funding of the Pavlov Institute of Physiology, Russian Academy of Sciences № 124020100105-6 (for O.G.).

## Notes

https://github.com/DmitriiPerevozniuk/DRG_Cross_Excitation

### Competing Interest Statement

The authors have declared no competing interest.

